# Kinetochores respond to subtle changes in the stability of microtubule attachments

**DOI:** 10.1101/2021.02.19.432040

**Authors:** Jessica D. Warren, Sarah Y. Valles, Duane A. Compton

## Abstract

Proper attachment of spindle microtubules to kinetochores is necessary to satisfy the spindle assembly checkpoint and ensure faithful chromosome segregation. Microtubules detach from kinetochores to correct improperly oriented attachments, and overall kinetochore-microtubule (k-MT) attachment stability is determined in response to regulatory enzymes and the activities of kinetochore-associated microtubule stabilizing and destabilizing proteins. However, it is unknown whether regulatory enzyme activity or kinetochore-associated protein localization respond to subtle changes in k-MT attachment stability. To test for this feedback response, we monitored Aurora B kinase activity and the localization of select kinetochore proteins in metaphase cells following treatments that subtly stabilize or destabilize k-MT attachments using low dose Taxol or UMK57 (an MCAK agonist), respectively. Increasing k-MT stability induced changes in the abundance of some kinetochore proteins. In contrast, reducing k-MT stability induced both increases in Aurora B kinase signaling and changes in the abundance of some kinetochore proteins. Thus, kinetochores dynamically respond to changes in the stability of their attached microtubules. This feedback control contributes to tuning k-MT attachment stability required for efficient error correction to facilitate faithful chromosome segregation.

**Summary Statement:** Live cell imaging demonstrates that kinetochore signaling responds to feedback from attached microtubules to tune their stability to ensure faithful chromosome segregation during cell division.

## Introduction

Kinetochore-microtubule (k-MT) attachment stability systematically increases during the course of mitosis to ensure timely mitotic progression and high chromosome segregation fidelity (Bakhoum et al., 2009; Bakhoum and Compton, 2012; Kabeche and Compton, 2013; Zhai et al., 1995). Changes in k-MT attachment stability are generated by mitotic kinases [e.g. CDK1 (Dumitru et al., 2017; Kabeche and Compton, 2013; Vasquez et al., 1997), Plk1 (Hood et al., 2012; Liu et al., 2012), Aurora A/B (Cimini et al., 2006; Liu and Lampson, 2009; Ye et al., 2015)] and phosphatases [e.g. PP1 (Liu et al., 2010) and PP2A (Meppelink et al., 2015)] that modify the localization, activities, and/or microtubule-binding affinities of the conserved KMN network (KNL1–MIS12– NDC80 complexes) (Cheeseman et al., 2006; DeLuca et al., 2005; Tanaka, 2012) and a bevy of other proteins that further stabilize or destabilize k-MT attachments (DeLuca et al., 2011; Welburn et al., 2010; Zaytsev et al., 2014). In this well-established view, the stability of k-MT attachments is responding to, and is determined by, the activities of kinetochore proteins.

It has also been demonstrated that kinetochore composition changes if k-MT attachments are removed using high doses of microtubule targeting drugs. For example, the spindle assembly checkpoint protein Mad2 is displaced from kinetochores upon microtubule binding, but returns if those microtubule attachments are subsequently destroyed with high doses of nocodazole (Cassimeris et al., 1990; Magidson et al., 2016; Wan et al., 2009). Other studies have demonstrated significant changes in kinetochore architecture in response to gain or loss of k-MT attachment (Magidson et al., 2015; Sacristan et al., 2018; Waters et al., 1998). However, the sensitivity for this type of feedback remains unknown. For example, once k-MT attachments are established, does merely altering attachment stability (as opposed to the loss of attachment) elicit feedback that alters the localization and/or activity of kinetochore proteins?

To test for feedback under these conditions, we induced subtle changes in k-MT attachment stability in metaphase cells and monitored kinetochore protein localization and the activity of Aurora B kinase using a FRET-based biosensor to measure localized phosphorylation. Subtle stabilization of k-MT attachments reduced average inter-kinetochore distance (IKD) and increased Astrin kinetochore localization, but did not significantly alter Aurora B kinase activity. In contrast, subtle destabilization of k-MT attachments did not change IKD, but increased kinetochore localization of Hec1 and Astrin, decreased kinetochore localization of HURP, and increased Aurora B kinase activity. Importantly, most changes induced by destabilizing k-MT attachments are reversed by subsequent stabilization of k-MT attachments. These data reveal that kinetochore composition and activity dynamically respond to changes in the stability of k-MT attachments, and that feedback mechanisms are responsive to changes in k-MT stability of as little as ~23%.

## Results and Discussion

To monitor Aurora B activity at kinetochores in live cells, we generated a stable U2OS cell line expressing a previously characterized, Mis12-targeted, Aurora B-dependent FRET biosensor (Liu and Lampson, 2009). In the presence of nocodazole, the kinetochore-localized FRET emission ratio significantly increased (i.e. phosphorylation decreased) in U2OS cells upon treatment with the Aurora B kinase-specific inhibitor, ZM447439, consistent with the response previously published in mitotic HeLa cells (Liu and Lampson, 2009) (Figures S1A and S1B). The FRET emission ratio also significantly increased in the absence of nocodazole, when prometaphase cells were treated with ZM447439 (Figure S1C). This increase was not observed for metaphase cells, suggesting that the biosensor is maximally dephosphorylated in metaphase (Figure S1C). Furthermore, the kinetochore-localized FRET emission ratio significantly decreased (i.e. phosphorylation increased) upon treatment with the phosphatase inhibitor okadaic acid and was restored to control levels when cells were simultaneously treated with the kinase and phosphatase inhibitors (Figure S1C). These data indicate that the extent of phosphorylation of the biosensor reflects ongoing kinase (phosphorylation) and phosphatase (dephosphorylation) activities.

Next, we used the FRET biosensor to simultaneously measure inter-kinetochore distance (IKD) and the level of Aurora B kinase activity on individual kinetochores. The FRET emission ratio tended to increase (i.e. phosphorylation decreased) on the population of kinetochores as cells transit from prometaphase to metaphase (Figure S1D), indicating reduced Aurora B kinase activity relative to phosphatase activity. The average inter-kinetochore distance (IKD) also tended to increase on the population of kinetochores as cells transit from prometaphase to metaphase (Figure S1E). However, this population view masks substantial variation in phosphorylation between individual kinetochores (Figure 1). IKD is variable from chromosome to chromosome in metaphase cells at any given moment as kinetochores ‘breath’ (i.e. transiently converge and diverge from each other over time) during chromosome oscillatory movement (He et al., 2000; Jaqaman et al., 2010). Surprisingly, there is no significant correlation between FRET emission ratio on individual metaphase kinetochores relative to IKD (Pearson’s correlation, r = 0.03, p = 0.668) over the complete range of IKDs measured (Figure 1A). Similarly, the differential of FRET emission ratio between sister kinetochores within pairs was not correlated with IKD (Pearson’s correlation, r = 0.08, p = 0.476) over the complete range of IKDs measured (Figure 1B). Furthermore, the FRET emission ratio on individual kinetochores is uncoordinated between sisters in prometaphase (Pearson’s correlation, r = 0.07, p = 0.599) and displays moderate positive correlation in metaphase (Pearson’s correlation, r = 0.40, p = 0.002) (Figures 1C and 1D). These data indicate a degree of kinetochore autonomy in the net phosphorylation of an Aurora B kinase substrate localized via Mis12.

**Figure 1.**
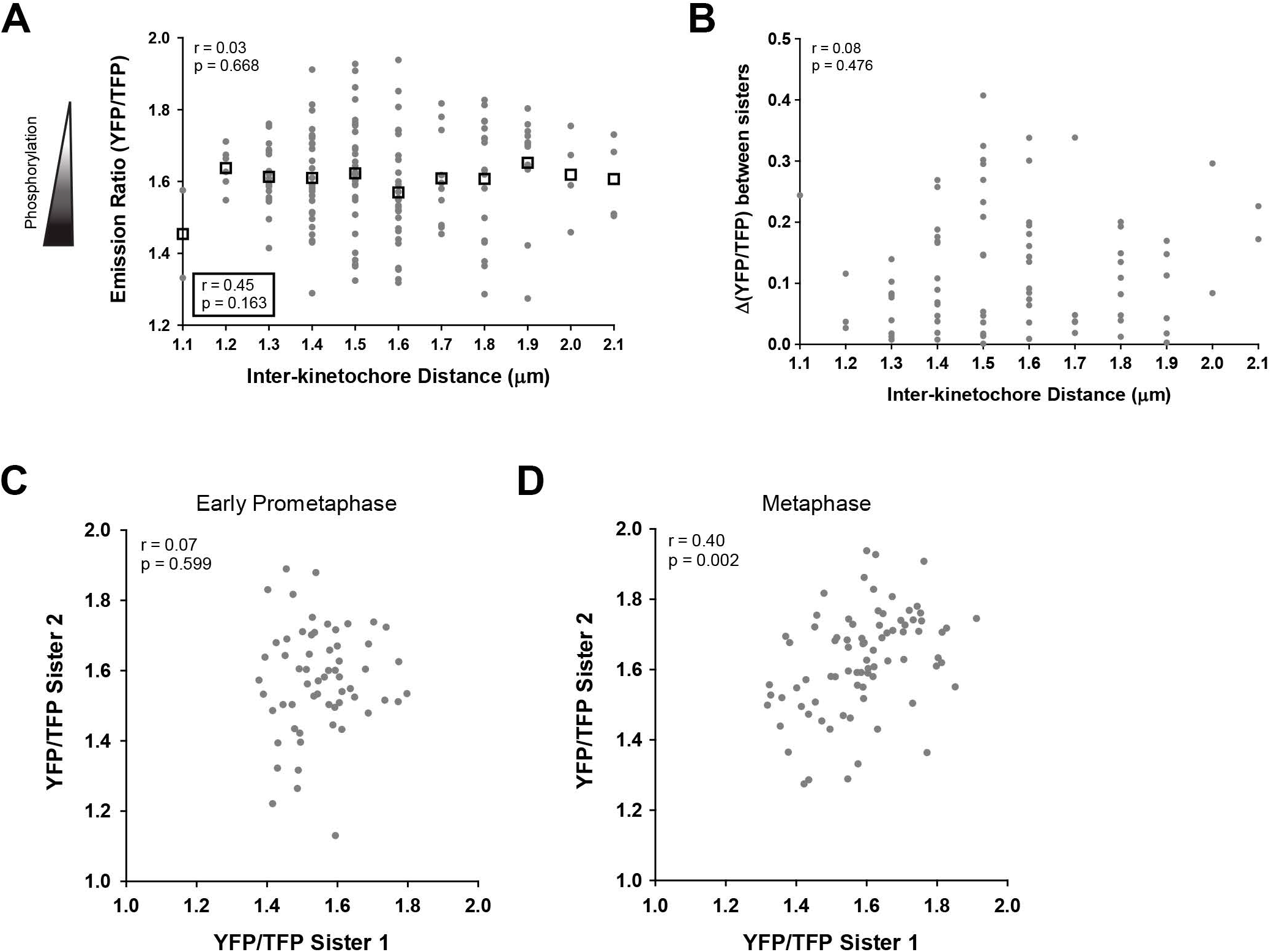
Phosphorylation of an Aurora B Mis12-targeted FRET sensor is not correlated with IKD in metaphase cells. **A)** Emission ratio data points (gray) for the metaphase cells quantified in Figure S1D plotted vs. IKD. IKD was measured as the distance between Mis12 localized YFP foci. For any given IKD, phosphorylation of the Mis12-targeted sensor is highly variable (n = 160 kinetochores) and is not correlated with IKD. Boxes represent the average emission ratio for each IKD. Correlation for individual data points (top) and IKD averages (bottom, boxed) was measured by the Pearson correlation co-efficient (r) and was considered significant if p < 0.05. **B)** The absolute difference between the emission ratios of sister kinetochores (Δ(YFP/TFP)) from (A) is plotted vs. IKD. Δ(YFP/TFP) varies over a range of IKDs and is not correlated with IKD. Correlation was measured by the Pearson correlation co-efficient (r) and was considered significant if p < 0.05. **C)** Emission ratio scatter plot for the early prometaphase and **D)** metaphase cells quantified in Figure S1D, where each gray spot represents a pair of sister kinetochores and their corresponding emission ratio (YFP/TFP) values. Aurora B dependent phosphorylation of sister kinetochores is uncoordinated in early prometaphase and becomes coordinated in metaphase. Correlation was measured by the Pearson correlation co-efficient (r) and was considered significant if p < 0.05.

To test whether kinetochores have the capacity to respond to changes in k-MT attachment stability, we treated U2OS cells with nanomolar concentrations of Taxol or UMK57 to decrease or increase the rate of microtubule detachment from kinetochores, respectively. We limited these analyses to cells in metaphase to prevent any potentially confounding effects related to the prometaphase to metaphase transition, and to ensure that chromosomes had established robust kinetochore-microtubule attachments. Cells treated with 5 nM Taxol for 1 hour were delayed slightly in mitotic completion and most cells progressed to anaphase albeit with a greater proportion of multipolar spindles (Figures S2A and S2B), consistent with previous studies (Brito and Rieder, 2009; Zasadil et al., 2014). The slight delay in mitotic completion induced by Taxol treatment was not accompanied by a significant increase in the percentage of metaphase cells with detectable Mad2-positive kinetochores (Figure S2C).

The metaphase k-MT detachment rate, as determined by measuring fluorescence dissipation of a photoactivated spindle region (Kabeche and Compton, 2013), significantly decreased in the presence of 5 nM Taxol such that the overall k-MT half-life increased by ~26% compared to DMSO-treated cells (Figures 2A and 2B). Taxol treatment also reduced metaphase IKD by ~9% (Figure 2C). However, under these conditions there was only a slight increase in phosphorylation at the location of the Mis12-targeted FRET sensor which was not significant (Figure 2D).

**Figure 2.**
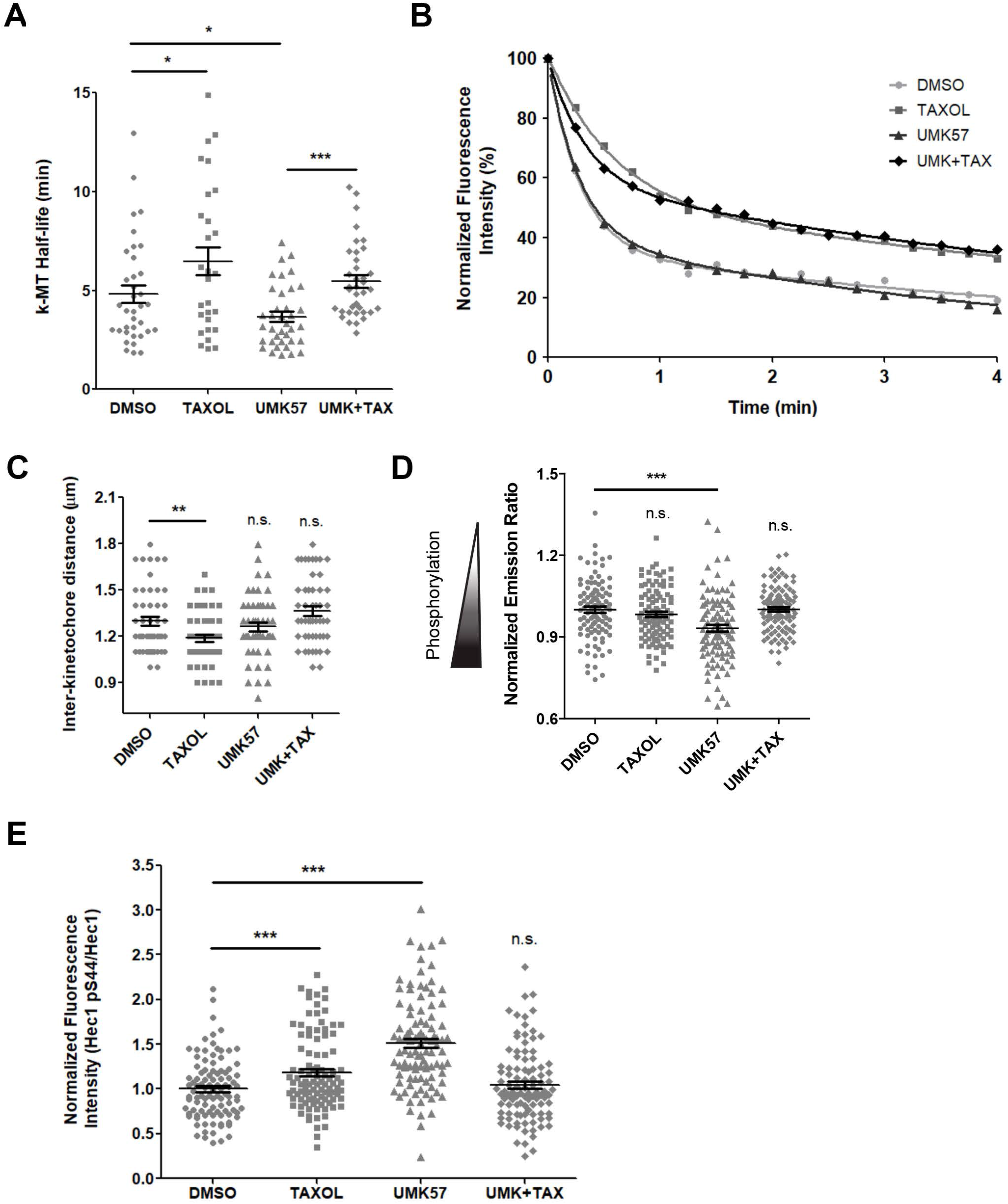
Aurora B phosphorylation increases in response to the destabilization of k-MT attachments. **A)** Average k-MT half-life for DMSO (circles), Taxol (squares), UMK57 (triangles), or UMK57+Taxol (diamonds)-treated U2OS cells expressing photoactivatable GFP-α-tubulin. Error bars indicate SEM; n ≥ 28 metaphase cells per condition; *p ≤ 0.05; ***p ≤ 0.001 using two-tailed t test. **B)** Examples of normalized fluorescence dissipation after photoactivation in metaphase cells treated with DMSO (light gray circles), Taxol (gray squares), UMK57 (dark gray triangles), or UMK57+Taxol (black diamonds). Decay curves are from cells that are representative of the average k-MT half-life in (A). **C)** U2OS cells stably expressing a Mis12-targeted Aurora B FRET sensor were treated for 1 hour with DMSO (circles), Taxol (squares), UMK57 (triangles), or UMK57+Taxol (diamonds) in the presence of 5 μM MG-132 and imaged to measure inter-kinetochore distance (IKD). Error bars indicate SEM; n = 100 kinetochores per condition; **p ≤ 0.01; n.s., p ≥ 0.05 using two-tailed t test. **D)** Corresponding YFP/TFP emission ratio measurements for kinetochore pairs quantified in (C). The decreased emission ratio observed with UMK57 treatment indicates increased phosphorylation of the FRET sensor. Error bars indicate SEM; n = 100 kinetochores per condition; ***p ≤ 0.001; n.s., p ≥ 0.05 using two-tailed t test. **E)** Quantification of Hec1 pS44 fluorescence intensity relative to total Hec1 fluorescence intensity. U2OS cells treated for 3 hours with DMSO (circles), Taxol (squares), UMK57 (triangles), or UMK57+Taxol (diamonds). Error bars indicate SEM; n = 100 kinetochores per condition; ***p ≤ 0.001; n.s., p ≥ 0.05 using two-tailed t test.

Complementary immunofluorescence analysis in fixed cells for phosphorylated Hec1 (pS44) demonstrated a significant increase in phosphorylation, suggesting that outer kinetochores may be more sensitive to increases in Aurora B activity (or decreases in phosphatase activity) (Figure 2E). With respect to protein localization, treatment of metaphase cells with low dose Taxol resulted in a significant (~33%) increase in the quantity of Astrin at kinetochores (Figures 3A and 3B). This change in protein localization is likely to be specific, since the localization of Hec1, HURP, and Ska3 did not significantly change with this dose of Taxol treatment (Figures 3C, 3D, and 4A-D).

**Figure 3.**
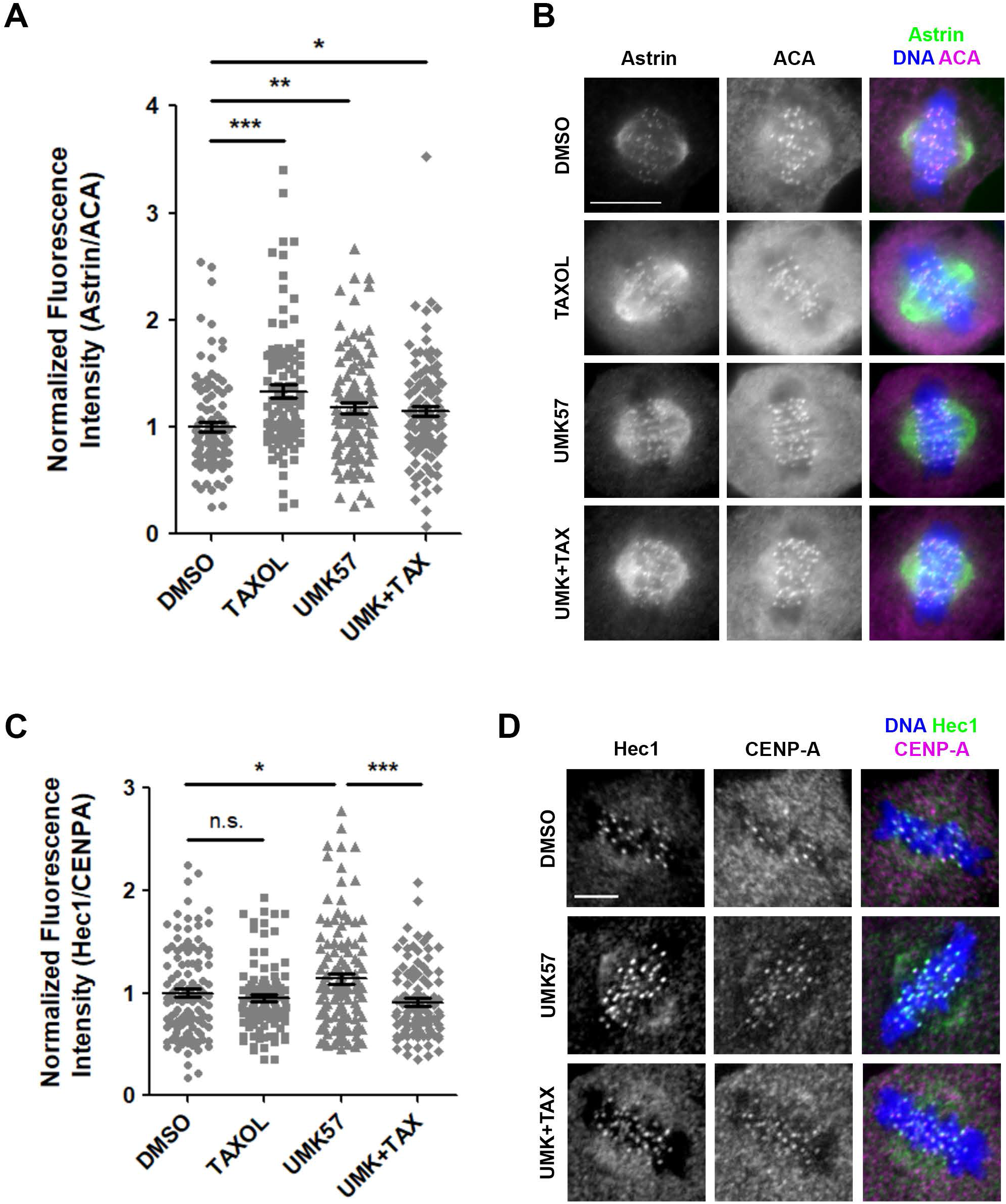
k-MT destabilization increases Astrin and Hec1 kinetochore localization. **A)** Quantification of Astrin fluorescence intensity relative to ACA. U2OS cells treated for 3 hours with DMSO (circles), Taxol (squares), UMK57 (triangles), or UMK57+Taxol (diamonds). Error bars indicate SEM; n = 100 kinetochores per condition; *p ≤ 0.05; **p ≤ 0.01; ***p ≤ 0.001 using two-tailed t test. **B)** Representative images of U2OS cells treated for 3 hours with DMSO, Taxol, UMK57, or UMK57+Taxol and then fixed and stained for DNA (blue), Astrin (green), and ACA (magenta). Images represent single slices of fluorescence z-stacks. Scale bar, 10 μm. **C)** Quantification of Hec1 fluorescence intensity relative to CENP-A. U2OS cells treated for 3 hours with DMSO (circles), Taxol (squares), UMK57 (triangles), or UMK57+Taxol (diamonds). Error bars indicate SEM; n ≥ 100 kinetochores per condition; *p ≤ 0.05; ***p ≤ 0.001; n.s., p ≥ 0.05 using two-tailed t test. **D)** Representative images of U2OS cells treated for 3 hours with DMSO, UMK57, or UMK57+Taxol and then fixed and stained for DNA (blue), Hec1 (green), and CENP-A (magenta). Images represent single slices of fluorescence z-stacks. Scale bar, 5 μm.

**Figure 4.**
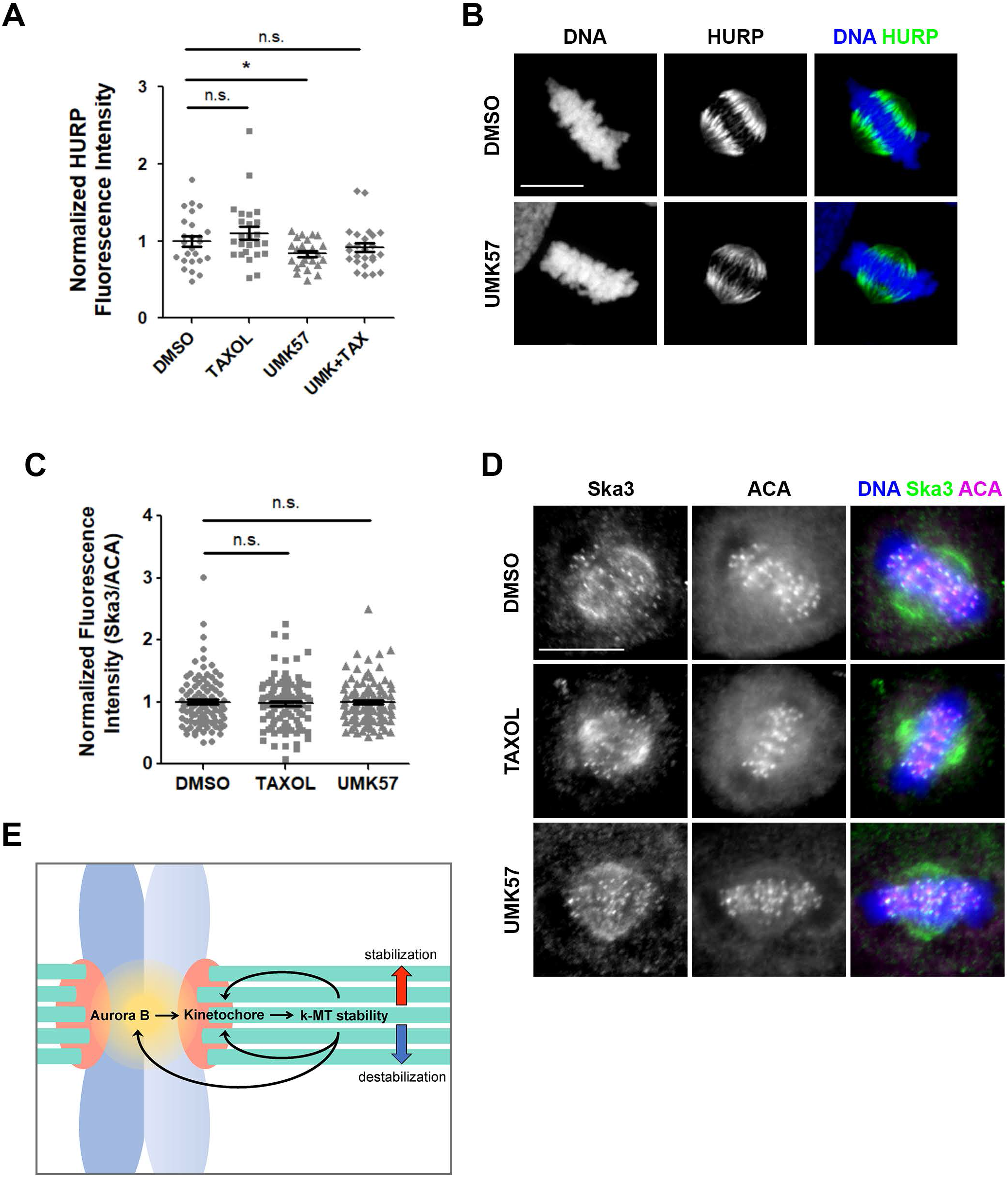
k-MT destabilization decreases HURP spindle localization, while Ska3 kinetochore localization is unchanged. **A)** Quantification of HURP fluorescence intensity. Whole spindle measurements were made on sum intensity projections of fluorescence z-stacks. U2OS cells treated for 3 hours with DMSO (circles), Taxol (squares), UMK57 (triangles), or UMK57+Taxol (diamonds). Error bars indicate SEM; n = 25 spindles per condition; *p ≤ 0.05; n.s., p ≥ 0.05 using two-tailed t test. **B)** Representative images of U2OS cells treated for 3 hours with DMSO or UMK57 and then fixed and stained for DNA (blue) or HURP (green). Images represent maximum intensity projections of fluorescence z-stacks. Scale bar, 10 μm. **C)** Quantification of Ska3 fluorescence intensity relative to ACA. U2OS cells treated for 3 hours with DMSO (circles), Taxol (squares), or UMK57 (triangles). Error bars indicate SEM; n = 100 kinetochores per condition; n.s., p ≥ 0.05 using two-tailed t test. **D)** Representative images of U2OS cells treated for 3 hours with DMSO, Taxol, or UMK57 and then fixed and stained for DNA (blue), Ska3 (green), and ACA (magenta). Images represent single slices of fluorescence z-stacks. Scale bar, 10 μm. **E)** A proposed model for the regulation of k-MT attachment stability. Aurora B influences kinetochore substrates to determine k-MT stability. There is feedback from k-MT attachment sites in response to k-MT stabilization or destabilization. This feedback may influence kinetochore protein composition directly or indirectly (through the modulation of Aurora B activity).

Next, we increased microtubule detachment rates from kinetochores in metaphase cells using 100 nM UMK57, a previously characterized MCAK agonist (Orr et al., 2016). Cells treated with UMK57 for 1 hour displayed no delay in mitotic completion or progression to anaphase and no change in the percentage of detectable Mad2-positive metaphase cells, and no change in the progression of cells to anaphase (Figures S2A-C) (Orr et al., 2016). The k-MT detachment rate, as determined by measuring fluorescence dissipation of a photoactivated spindle region (Kabeche and Compton, 2013), increased in the presence of 100 nM UMK57 such that the overall k-MT half-life decreased by ~23% compared to DMSO-treated cells (Figures 2A and 2B). In addition, the kinetochore-localized FRET emission ratio was significantly reduced (i.e. phosphorylation increased) in metaphase cells treated with UMK57 (Figure 2D). This is consistent with the previously reported increase in the quantity of active Aurora B kinase as judged by increased levels of phosphorylated T232 at centromeres (Orr et al., 2016). The observed increase in phosphorylation of a kinetochore-localized Aurora B substrate was further confirmed by immunofluorescence using a Hec1 phospho-specific antibody. The amount of phosphorylated Hec1 (pS44) at kinetochores significantly increased in response to k-MT destabilization (Figure 2E). Interestingly, IKD did not significantly change following UMK57 treatment (Figure 2C), indicating that the observed changes in phosphorylation are independent of IKD. UMK57 treatment significantly increased the localization of Astrin (~18%; Figures 3A and 3B) and Hec1 (~14%; Figures 3C and 3D) to kinetochores, and decreased the localization of HURP (Silljé et al., 2006) to spindle microtubules (~16%; Figures 4A and 4B). These changes in protein localization appear to be specific because there was no change in the localization of Ska3 protein to kinetochores in UMK57 treated cells (Figures 4C and 4D).

The changes induced at metaphase kinetochores by treatment with UMK57 could result from either the alteration of MCAK activity or the destabilization of k-MT attachments. To distinguish between these possibilities, we tested if the kinetochore changes that we observed in response to UMK57 treatment could be reversed by the subsequent stabilization of k-MT attachments with low dose Taxol, which specifically targets microtubules. For these analyses, we treated cells with 100 nM UMK57 for 30 minutes and subsequently added 5 nM Taxol for an additional 30 minutes. Previous *in vitro* studies confirmed that UMK57 retains the capacity to increase MCAK activity on Taxol-stabilized microtubules (Orr et al., 2016). The detachment rate of microtubules from kinetochores in metaphase cells treated with the combination of UMK57 and Taxol was significantly decreased relative to cells treated with UMK57 alone. The k-MT half-life for UMK57+Taxol-treated cells increased by ~33% (relative to UMK57 alone) and was no longer significantly different from that of control cells (Figures 2A and 2B). Moreover, the reduction in IKD induced by Taxol treatment alone was prevented if cells were initially treated with UMK57 (Figure 2C). Furthermore, the FRET emission ratio in metaphase cells treated with the combination of UMK57 and Taxol was significantly increased (e.g. phosphorylation decreased) relative to cells treated with UMK57 alone and was no longer significantly different from that of control cells (Figures 2D). The observed change in phosphorylation of a kinetochore-localized Aurora B substrate was further confirmed by immunofluorescence using a Hec1 (pS44) phospho-specific antibody (Figure 2E). The changes in kinetochore localization of both Hec1 and HURP induced by UMK57 were reversed by subsequent treatment with low dose Taxol and were no longer significantly different from control cells (Figures 3C and 4A), while Astrin localization remained elevated under each experimental condition (Figure 3A). These data indicate that kinetochore protein localization and/or activity change in response to altered k-MT attachment stability.

It has been demonstrated that kinetochores have the capacity to respond to extreme changes in k-MT attachment status. An example is the re-recruitment of Mad2 back to kinetochores following the obliteration of all k-MT attachments (Waters et al., 1998) using concentrations of nocodazole that completely depolymerize all kinetochore microtubules (Cassimeris et al., 1990). Such extreme treatments are, arguably, not physiological, which is why we sought to test if kinetochores respond to more subtle changes in k-MT attachment stability. Under these conditions, we demonstrate that kinetochore composition and the phosphorylation status of an Aurora B kinase substrate respond rapidly to physiologically relevant changes in k-MT attachment stability, since the extent of k-MT attachment stability changes induced here are within the dynamic range of changes that occur between different stages of normal mitosis (Kabeche and Compton, 2013). Whereas the magnitude of change in phosphorylation observed under these conditions appears small, in companion work we recently demonstrated that Hec1 is only partially phosphorylated (regardless of MT attachment status at the kinetochore) and that k-MT attachment stability is highly sensitive to very small changes in the extent of Hec1 phosphorylation (Kucharski, Hards, Godek, Gerber, & Compton; under review). Thus, the degree of phosphorylation change detected also lands within a physiologically relevant dynamic range. Remarkably, the responses we observe are independent of changes in inter-kinetochore distance, indicating that in metaphase human cells, as has been shown in other cell types, that spatial distance relative to the centromere is not the sole determinant regulating the extent of phosphorylation of Aurora B kinase kinetochore substrates (Broad et al., 2020; Campbell and Desai, 2013; Godek et al., 2015; Hadders et al., 2020; Liu et al., 2010). On the contrary, our results reveal that kinetochore protein localization and the extent of phosphorylation of an Aurora B kinase substrate are responsive to the stability of k-MT attachments, which reveals the presence of feedback mechanisms from sites of microtubule attachment (or occupancy) to the kinetochore. In this context, Aurora B kinase promotes the destabilization of k-MT attachments, but there is feedback from less stable k-MT attachments to influence kinetochore composition and Aurora B kinase to adjust their effect. Such feedback would contribute to establishing and maintaining k-MT attachment stability at specific “set points” during different phases of mitosis and is consistent with the principles of homeostatic regulation (Figure 4E).

Understanding how feedback participates in specifying k-MT attachment stability at different mitotic phases, why we observe a selective response of kinetochore constituents to either k-MT stabilization or destabilization, or how kinetochores shift from acting autonomously in prometaphase to more coordination between sisters in metaphase must await a more fulsome understanding of the molecular constituents of the signaling network that is responding to k-MT attachment stability at kinetochores. In this context, we propose a working model where a signal emanates from the site of microtubule plus end attachment within kinetochores, and that a signal transduction pathway including protein kinases and phosphatases responds to microtubule end dwell time at the point of attachment. Since changes in k-MT detachment rate will necessarily translate into changes in microtubule occupancy at the kinetochore, it is possible that microtubule occupancy provides the primary signal in this context, similar to its established role in SAC signaling (Etemad et al., 2019; Kuhn and Dumont, 2019).

## Acknowledgments

We thank all members of our laboratory for stimulating discussions and critical feedback in the interpretation and analysis of results. We also thank Benjamin Kwok, Alexey Khodjakov, Michael Lampson, Jennifer DeLuca, and Erich Nigg for kindly providing important materials and reagents. This work was supported by grant R37GM051542 from the National Institutes of Health (D.A.C.).

## Author Contributions

Conceptualization, J.D.W. and D.A.C.; Methodology, J.D.W., S.Y.V. and D.A.C.; Formal Analysis, J.D.W., S.Y.V.; Writing, J.D.W., S.Y.V. and D.A.C.; Visualization, J.D.W. and D.A.C.; Supervision, D.A.C.; Funding Acquisition, D.A.C.

## Supplementary Figure Legends

**Figure S1.**
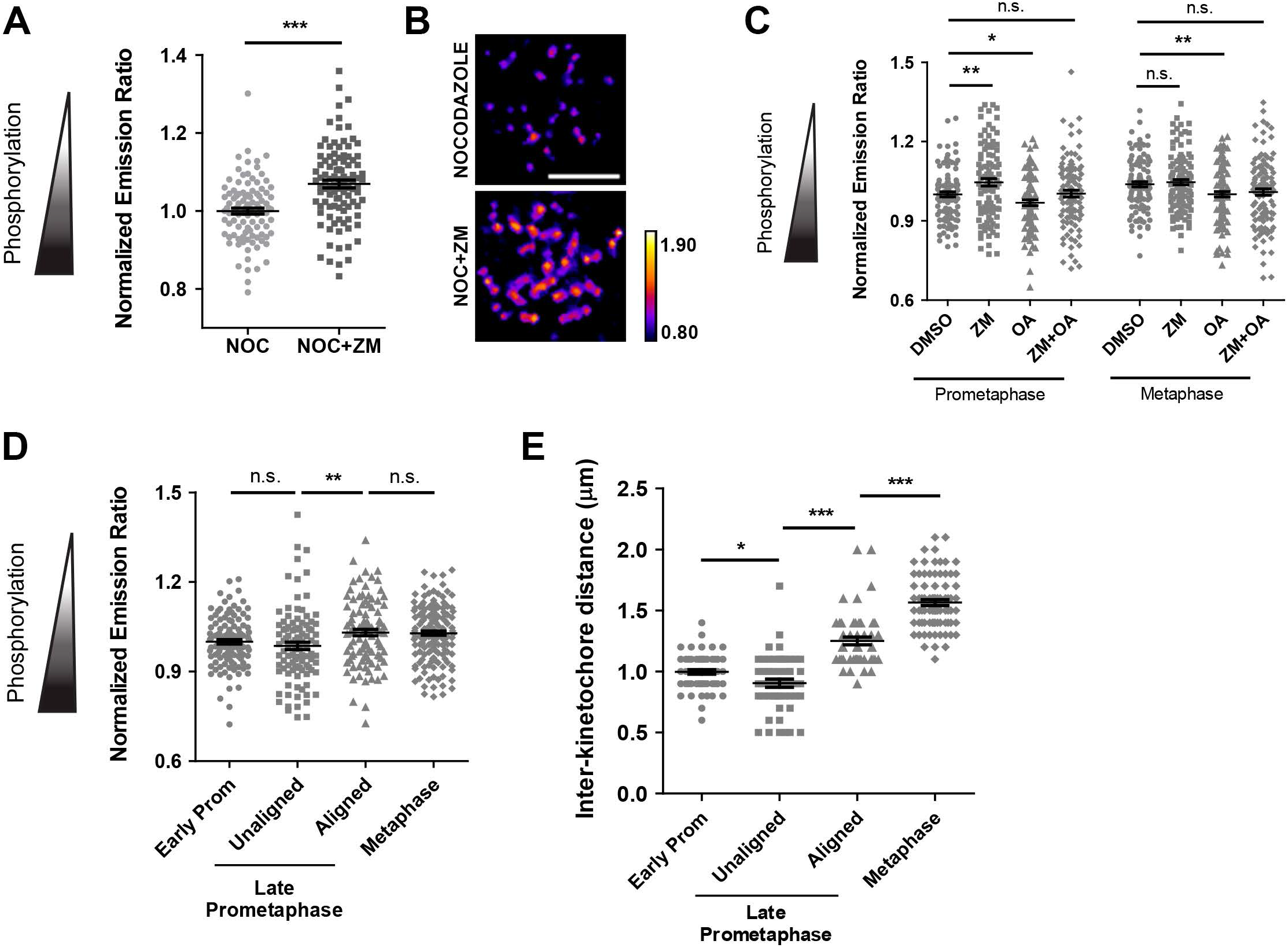
A Mis12-targeted Aurora B FRET sensor serves a readout for Aurora B activity during mitosis. **A)** U2OS cells stably expressing Mis12-targeted Aurora B FRET sensor treated for 1 hour with nocodazole (NOC) or nocodazole + ZM447439 (NOC+ZM). The YFP/TFP emission ratio increases in cells treated with the Aurora kinase inhibitor, indicating sensor dephosphorylation. Error bars indicate SEM; n = 100 kinetochores from N = 10 cells per condition; ***p ≤ 0.001 using two-tailed t test. **B)** Representative images of YFP/TFP emission ratio at kinetochores quantified in (A). Images represent single slices of fluorescence z-stacks. Scale bar, 5 μm. **C)** U2OS cells stably expressing Mis12-targeted Aurora B FRET sensor were imaged in prometaphase or metaphase to calculate YFP/TFP ratio. Cells were treated for 30 minutes with MG132 before the addition of DMSO, ZM447439 (ZM), Okadaic Acid (OA) for 1 hour, or ZM+OA for 30 minutes + 30 minutes (added sequentially). Lower values indicate increased phosphorylation of the FRET sensor. Each treatment condition is compared to the DMSO treatment for the corresponding mitotic stage. Error bars indicate SEM; n ≥ 100 kinetochores analyzed per condition; *p ≤ 0.05; **p ≤ 0.01; n.s., p ≥ 0.05 using two-tailed t test. **D)** U2OS cells stably expressing Mis12-targeted Aurora B FRET sensor were imaged at various mitotic stages to calculate YFP/TFP ratio. 5 μM MG-132 was added prior to rose chamber assembly. Cells classified based on chromosome alignment as early prometaphase (Early Prom), late prometaphase, or metaphase. Late prometaphase kinetochore pairs were further classified as either unaligned or aligned. Lower values indicate increased phosphorylation of the FRET sensor. Error bars indicate SEM; n ≥ 102 kinetochores analyzed at each mitotic stage; **p ≤ 0.01; n.s., p ≥ 0.05 using two-tailed t test. **E)** Corresponding inter-kinetochore distance (IKD) measurements for cells quantified in (D). Error bars indicate SEM; n ≥ 102 kinetochores analyzed at each mitotic stage; *p ≤ 0.05; **p ≤ 0.01; ***p ≤ 0.001 using two-tailed t test.

**Figure S2.**
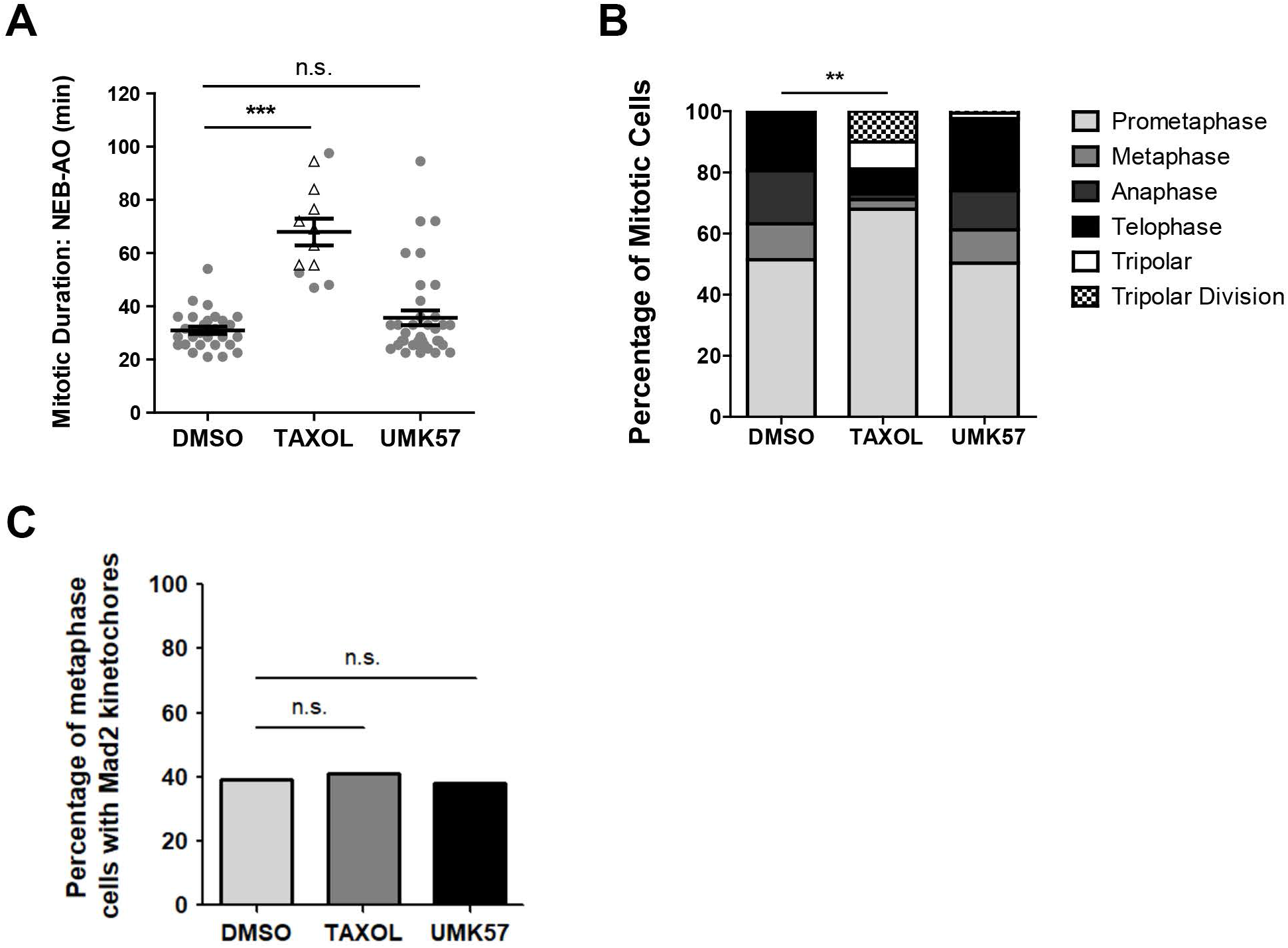
Effects of Taxol and UMK57 treatment on mitotic progression and the percentage of Mad2-positive kinetochores in metaphase. **A)** Mitotic duration measured as the time from nuclear envelope breakdown (NEB) to anaphase onset (AO) by DIC live cell imaging. U2OS cells treated with DMSO, Taxol, or UMK57 for 1 hour prior to the start of image acquisition. Each spot represents a single cell with a bipolar (gray circle) or tripolar (white triangle) spindle; bars indicate the mean ± SEM; n ≥ 12 cells per condition; n.s., p ≥ 0.05; ***p ≤ 0.001 using two-tailed t test. **B)** Classification of mitotic stages observed in fixed U2OS cells treated for 3 hours with DMSO, Taxol, or UMK57. The percentage of prometaphase cells is significantly increased with Taxol treatment. n ≥ 96 cells per condition; **p ≤ 0.01 using two-tailed Fisher’s Exact Test. **C)** Percentage of metaphase cells with at least one Mad2-positive kinetochore. U2OS cells treated for 3 hours with DMSO, Taxol, or UMK57. n = 122 metaphase cells per condition; n.s., p ≥ 0.05 using two-tailed Fisher’s Exact Test.

## Methods

### Drug treatment

UMK57 was used at 100 nM, as previously reported (Orr et al., 2016). Paclitaxel (Taxol) was used at 5 nM (Biotang). The following drugs were also used at the specified concentrations: MG-132 (5 μM; Sigma-Aldrich), nocodazole (100 ng/mL; Sigma Aldrich), ZM447439 (3 μM; Tocris Bioscience), Okadaic Acid (100 nM; Adipogen Life Sciences). All controls for drug treatment were performed using 0.1% DMSO.

### Cell culture

U2OS (ATCC®, HTB-96) cells were grown in Dulbecco’s modified Eagle’s medium, (DMEM; Invitrogen). U2OS cells expressing the Mis12-targeted Aurora B FRET sensor (Addgene, #45231) or photoactivatable GFP-α-tubulin (plasmid provided by Alexey Khodjakov) were cultured in DMEM supplemented with 1 mg/mL G418 (InvivoGen). All cell lines were grown in medium supplemented with 10% FBS (HyClone), 10 mM HEPES (GE Healthcare), 250 μg/L Amphotericin B (Sigma Aldrich), 50 U/mL penicillin (Mediatech) and 50 μg/mL streptomycin (Mediatech). All cell lines were grown at 37°C in a humidified atmosphere with 5% CO2. All live cell imaging was performed in FluoroBrite DMEM (Gibco) supplemented with 10% FBS, 4mM L-glutamine (Thermo Fisher Scientific), 10 mM HEPES, 250 μg/L Amphotericin B, 50 U/mL penicillin and 50 μg/mL streptomycin. Plasmid transfections were performed using FuGENE 6 (Roche Diagnostics) following the manufacturer’s instructions.

### DIC live cell imaging

Live cell imaging was performed on a Nikon Eclipse Ti microscope with a Plan Apo VC 60x, 1.4 NA, oil immersion objective (Nikon). Cells were imaged in FluorBrite DMEM in a rose chamber maintained at 37°C by a heated stage. DIC z-stacks (1 μm step size, 12 μm range) were acquired with a cooled charge-coupled device camera (Andor Technology) every 1.5 minutes over the course of 16 hours.

### Photoactivation of spindle microtubules

A photoactivation assay was used to measure k-MT stability. U2OS cells stably expressing photoactivatable GFP-α-tubulin were maintained at 37°C in FluoroBrite DMEM during image acquisition. Cells were treated with DMSO, Taxol, or UMK57 for 1 hour prior to image acquisition and 5 μM MG-132 was added prior to rose chamber assembly to prevent mitotic exit. Images were acquired with a Plan Apo VC 100x, 1.4 NA, oil immersion objective (Nikon) using a QuorumWaveFX-X1 spinning disk confocal system on a Nikon Eclipse Ti microscope, equipped with an ILE laser source (Andor Technology), a Mosaic digital mirror (Andor Technology), and a Hamamatsu ImageEM camera. Metaphase cells were identified by differential interference contrast (DIC) microscopy. Photoactivation was performed with a 405 nm laser (35% power, 500ms pulse) on a rectangular region of interest over one half of the mitotic spindle, as previously described (Kabeche and Compton, 2013). Fluorescence z-stacks with a 1 μm step size (7 slices) were captured every 15 seconds for 4 minutes. Fluorescence dissipation after photoactivation (FDAPA) was quantified from maximum intensity projections using MetaMorph® software (Molecular Devices). Average pixel intensities were measured within an area surrounding the region of highest fluorescence intensity and background subtraction was performed using an equally sized area from the non-activated half-spindle at each time point. Fluorescence intensities were corrected for photobleaching using values of fluorescence loss obtained from photoactivated 1 μM Taxol stabilized spindles. Fluorescence intensities were then normalized to the first time point after photoactivation for each cell. To measure microtubule stability, the average fluorescence intensity at each time point was fit to a two-phase exponential decay curve 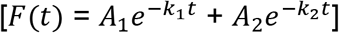 using GraphPad Prism software, where F(t) is measured photoactivated fluorescence at time t, A1 represents the percentage of photoactivated fluorescence from the fast decay process (non-k-MT population), and A2 represents the percentage of photoactivated fluorescence from the slow decay process (k-MT population), with decay rates of k1 and k2, respectively (Zhai et al., 1995). For high stringency, we only considered curves with good fit (R^2^ ≥ 0.98). The half-life (t^1/2^) for each process was calculated as ln2/k. The measured k-MT half-life represents the time it takes for half of the fluorescently labeled k-MTs to detach from kinetochores (Bakhoum et al., 2009).

### FRET sensor imaging and data analysis

Aurora B phosphorylation was measured in U2OS cells stably expressing a Mis12-targeted FRET biosensor, previously described in (Liu and Lampson, 2009). Live cells were imaged using a QuorumWaveFX-X1 spinning disk confocal system on a Nikon Eclipse Ti microscope, equipped with an ILE laser source (Andor Technology) and two Hamamatsu ImageEM cameras. TFP was excited at 445 nm, and TFP and YFP emissions were captured simultaneously with dual-camera acquisition. Fluorescence z-stacks with a 0.3 μm step size (21 slices) were acquired. Kinetochores were defined manually using Fiji (Schindelin et al., 2012) and the YFP/TFP emission ratio was calculated at each kinetochore. All FRET ratio results represent averages over multiple kinetochores from multiple cells. Pseudo-colored FRET images were generated using the Image Calculator function in Fiji. Inter-kinetochore distance was measured between the centroids of sister kinetochore (Mis12) spots located in the same focal plane.

### Antibodies

The following primary antibodies were used for immunofluorescence: ACA (anti-centromere antibody; Geisel School of Medicine; 1:1000), Astrin (C-terminal; 1:1000) (Mack and Compton, 2001), CENP-A (#2186; Cell Signaling; 1:500), Hec1 (C-11; Santa Cruz; 8 ug/mL), HURP (gift from Erich Nigg; 1:1000) (Silljé et al., 2006), Mad2 (PRB-452C; Covance; 1:500), pHec1 S44 (gift from Jennifer DeLuca; 1:500), Ska3 (gift from Erich Nigg; 1:1000) (Gaitanos et al., 2009). Secondary antibodies used were Alexa Fluor® 488, 568, 594, and 647 (Molecular Probes; 1:2000). DAPI (Molecular Probes) was used at 400 ng/mL.

### Immunofluorescence, fixed cell imaging, and data analysis

For visualization of Astrin, Hec1, HURP, and Ska3, cells were simultaneously permeabilized and fixed for 10 minutes at room temperature in PTEMF buffer (20 mM PIPES [pH 6.8], 4% formaldehyde, 0.2% Triton X-100, 10 mM EGTA, 1 mM MgCl2). Cells were washed with Tris-buffered saline with 5% bovine serum albumin (TBS-BSA) + 0.5% Triton X-100 for 2 × 10 minutes and then rinsed with TBS-BSA. Primary antibodies were diluted in TBS-BSA + 0.1% Triton X-100 and coverslips were incubated for 1-3 hours at room temperature. Cells were then washed with TBS-BSA + 0.1% Triton X-100 for 3 × 10 minutes with vigorous shaking. Secondary antibodies were diluted in TBS-BSA + 0.1% Triton X-100 + DAPI and coverslips incubated for 1-2 hours at room temperature. Cells were then washed with TBS-BSA + 0.1% Triton X-100 for 3 × 10 minutes with vigorous shaking and then mounted using ProLong® Gold antifade reagent (Molecular Probes). For visualization of Mad2, cells were treated prior to PFA fixation as follows: 1 minute MTSB [(pH 6.8); 4 M glycerol, 100 mM PIPES, 1 mM EGTA and 5 mM MgCl2], 2 minutes MTSB + 1.0% Triton X-100, 2 minutes MTSB. Antibody incubations and washes were then performed as previously described. Images were acquired with a cooled charge-coupled device camera (Andor Technology) mounted on a Nikon Eclipse Ti microscope with a Plan Apo VC 60x, 1.4 NA, oil immersion objective (Nikon). Image series in the z-axis were obtained using 0.2 μm optical sections (21 slices). Kinetochore fluorescence intensity (minus background) quantifications were performed on single Z-slices as previously described (Hoffman et al., 2001). Mean, whole-spindle HURP intensity (minus background) was measured on sum intensity projections of fluorescence z-stacks (0.5 μm step size; 17 slices).

For visualization of phosphorylated Hec1 (pHec1 S44), cells grown on coverslips were rinsed with pre-warmed PHEM buffer (60 mM PIPES, 25 mM HEPES, 10 mM EGTA, 4 mM MgSO4, at pH 6.9) before pre-extraction using a freshly made and sonicated lysis solution containing 2% Triton-X in 1x PHEM + 10mM glycerol 2-phosphate + 100nM microcystin for 7 minutes. Pre-warmed 1x PHEM+ 2% PFA was placed on coverslips and cells were fixed for 20 minutes at 37°C. Coverslips were then rinsed in 1x PHEM followed by three washes in 1x PHEM+ 0.1% Triton-X on an orbital shaker at high speed. Blocking was performed for 1 hour using 10% goat serum. Coverslips were incubated in primary antibody solution (5% goat serum in 1x PBS) at room temperature for 30 minutes on an orbital shaker at low speed and then overnight at 4°C. The following day, coverslips were washed three times in 1x PHEM+ 0.1% Triton-X on an orbital shaker at high speed and then rinsed in 1x PHEM. Coverslips were incubated for 1 hour in secondary antibody solution (5% goat serum in 1x PBS + were Alexa Fluor® 488 and 594; 1:300), away from light, on an orbital shaker at low speed. Coverslips were then washed three times in 1x PHEM+ 0.1% Triton-X on an orbital shaker at high speed followed by one rinse with 1x PHEM. DAPI was added to coverslips at 200ng/mL in 1x PHEM for 1 minute. Coverslips were then washed twice with 1x PHEM+ 0.1% Triton-X on an orbital shaker at high speed and once in 1x PHEM to remove residual DAPI solution. Coverslips were mounted on glass slides using ProLong® Gold (Molecular Probes). Images were acquired with an ORCA-Fusion Gen-III sCMOS camera (Hamamatsu) mounted on a Nikon Eclipse Ti microscope with a Plan Apo VC 100x, 1.4 NA, oil immersion objective (Nikon). Image series in the z-axis were obtained using 0.2 μm optical sections (21 slices). Kinetochore fluorescence intensity (minus background) quantifications were performed on single Z-slices as previously described (Hoffman et al., 2001).

### Statistical Analysis

All data analyzed as indicated in the figure legends. All graphical representations of data and statistical analyses were performed using GraphPad Prism software. Graphs present normalized data for clarity, but all statistical tests were performed on raw data values.

